# Valence-specific representation of uncertainty in the anterior cingulate is impaired in those with affective symptoms

**DOI:** 10.64898/2026.06.16.732356

**Authors:** Wanjun Lin, Laurence T. Hunt, Erdem Pulcu, Michael Browning

**Author notes:** co-senior authors (Erdem Pulcu, Michael Browning).

## Abstract

The ability to seek reward and avoid punishment is a fundamental survival instinct. In natural environments, however, the statistics of rewards and punishments can change independently of one another. In addition, individuals experiencing anxiety and depression may be selectively biased to process rewards and punishments differently. Here, we examine how humans adapt their behavior in such dynamic environments, using a task where cues are associated with independently changing probabilities of rewards (wins) and punishments (losses). We demonstrate that participants dynamically adjust their learning rates based on the relative volatility of each valence. Neuroimaging reveals that this behavioral flexibility is supported by valence-specific segregation of volatility signals within distinct subregions of the anterior cingulate cortex (ACC: perigenual and dorsal). Furthermore, individuals with higher levels of anxiety and depression exhibit a relative reduction in loss-volatility learning adaptation, accompanied by less distinct neural encoding of win and loss volatility in both ACC subregions. Together, our findings indicate that flexible adaptation to separate reward and punishment contingencies relies on distinct, valence-specific tracking of volatility within the prefrontal cortex. These findings suggest a potential computational and neuroanatomical framework for understanding maladaptive learning in affective disorders.

## Introduction

Both reward and punishment are effective reinforcers of human and animal behaviours (Marschark and Baenninger 2002, Sidin 2021). The “Carrot and stick” approach is commonly used in human education and animal training, where the general idea is to design rewards and punishments that point to the same desired behavioural outcomes. However, in naturalistic environments, reward and punishment might not favor the same actions. For example, approaching a riverside could be both rewarding (water) and punishing (predators such as crocodiles) for animals. In these situations, the probabilities of both rewards and punishments must be separately estimated to inform the required risk-benefit trade-off.

One factor that influences the impact of events on learning is uncertainty. Adaptive learning requires an ability to track different forms of uncertainty, including expected uncertainty (related to variability of outcomes) and unexpected uncertainty of the outcomes (related to variability of the environment) (Soltani and Izquierdo 2019). The latter, also known as volatility, particularly influences how rapidly an agent updates their belief about outcome probabilities following receipt of information (i.e. their learning rate). Specifically, studies have shown that people increase learning rates when the probabilities of reward are more volatile, i.e., when the contingencies of reward change more frequently (Behrens, Woolrich et al. 2007, Behrens, Hunt et al. 2008). Similar learning rate adjustments to volatility have also been observed in a punishment (electric-shock) learning task (Browning, Behrens et al. 2015). However, how people adapt their learning when both rewards and punishments are present is less studied. In previous work, we designed a novel task (the information bias learning task), where we presented participants with both rewards (monetary wins) and punishments (monetary losses), and manipulated the volatility of these two types of outcome independently across blocks. We showed that people adopt higher learning rates to the relevant valence domain i.e., to the one that was relatively more informative (i.e., volatile) than the other (Pulcu and Browning 2017). Here, we further develop the information bias learning task to examine the underlying neural mechanism using functional magnetic resonance imaging (fMRI).

The anterior cingulate cortex (ACC) is involved in processing reinforcement learning outcomes (both reward and punishment) (Holroyd and Yeung 2012), important for behavioral flexibility (Klein-Flügge, Bongioanni et al. 2022), and also implicated in the etiology of mood disorders (Drevets, Savitz et al. 2008, Price and Drevets 2012). In a simple probabilistic learning task, Monosov showed that distinct neuron populations in Macaques’ ACC encode outcomes and uncertainty for either rewards or punishments during associative learning (Monosov 2017). Similar results have shown that distinct neurons in the pregenual ACC encode positive and negative valence information when macaques make a decision to either approach or avoid a combination of appetitive and aversive outcomes (Amemori and Graybiel 2012). Human fMRI studies show that reward (monetary gain) increases BOLD signals in a more ventral region of ACC (rACC), while punishment (monetary loss) activates dorsal ACC (Fujiwara, Tobler et al. 2009). When outcome contingencies change over time (unexpected uncertainty), ACC responses to reward feedback increase as a function of volatility level during reward learning tasks in humans (Behrens, Woolrich et al. 2007, Behrens, Hunt et al. 2008, Silvetti, Seurinck et al. 2013) and non-human primates (Massi, Donahue et al. 2018). Here, we examine whether there is a distinct modulation effect of reward and punishment volatility on outcome signals in the ACC when both reward and punishment can be used to guide learning, and with independent volatility dynamics.

Environmental volatility has direct influences on human mental health. Real-world uncertainty, such as early life environmental unpredictability and beliefs in life’s unpredictability, are associated with anxiety and depression (Ross, Hood et al. 2016, Ross, Heming et al. 2023, Koss, Kronaizl et al. 2025). In experimental settings, people with higher depressive symptoms reported feeling less happy during reward learning in a volatile environment, but not in a stable environment (Blain and Rutledge 2020). Likewise, failing to adapt learning to punishment (electric shocks) in volatile environments was associated with acute emotional distress (De Berker, Rutledge et al. 2016). Consistent with this, we have reported an association between reduced punishment learning rate adaptation to volatility and trait anxiety, with no similar association found for rewarding outcomes (Browning, Behrens et al. 2015, Pulcu and Browning 2017). Gagne et al (Gagne, Zika et al. 2020) replicated the association for punishing outcomes with symptoms of both anxiety and depression, however they also reported poorer reward learning rate volatility adaptation using a separate reward learning task, suggesting a valence-general impairment in learning rate volatility adaptations in people with higher anxiety and depression. Here, we examine volatility adaptation when choices produce both rewarding and punishing outcomes in the same task, and test whether symptoms of anxiety and depression are associated with a differential adaptation to these two outcomes.

In the current study, non-clinical human participants, purposively recruited to have a range of depression symptoms, completed a learning task while fMRI data was collected. In the task, both reward (wins) and punishment (losses) outcomes were independently associated with choices and, critically, the informativity of wins and losses was manipulated by changing their volatility between blocks (Pulcu and Browning 2017). We show that participants make choices based on the relative informativity of wins and losses, that distinct ACC sub-regions (perigenual and dorsal ACC) respond to the volatility of reward and punishment volatility, respectively, and that people with higher anxiety and depression scores displayed a relatively reduced learning rate adaptation to losses, which was associated with less distinct win and loss volatility signals in ACC subregions, particularly a reduced loss volatility activity in the dorsal ACC.

## Results

We pre-screened non-clinical participants using a standard measure of depressive symptoms - the Quick Inventory of Depressive Symptoms (QIDS) (Rush, Trivedi et al. 2003) - and, from this group, invited 30 participants (14 females), selected to have a range of depression symptoms (from low to moderate levels), to complete the study. All participants completed a battery of symptom score questionnaires and performed the information bias learning task (see Fig. 1a&b and methods) in the MRI scanner. One participant was excluded from all analyses because they chose the same shape for all trials in one of the blocks (see Table. S 1 for a summary of the demographic details of the included participants).

**Fig. 1.**
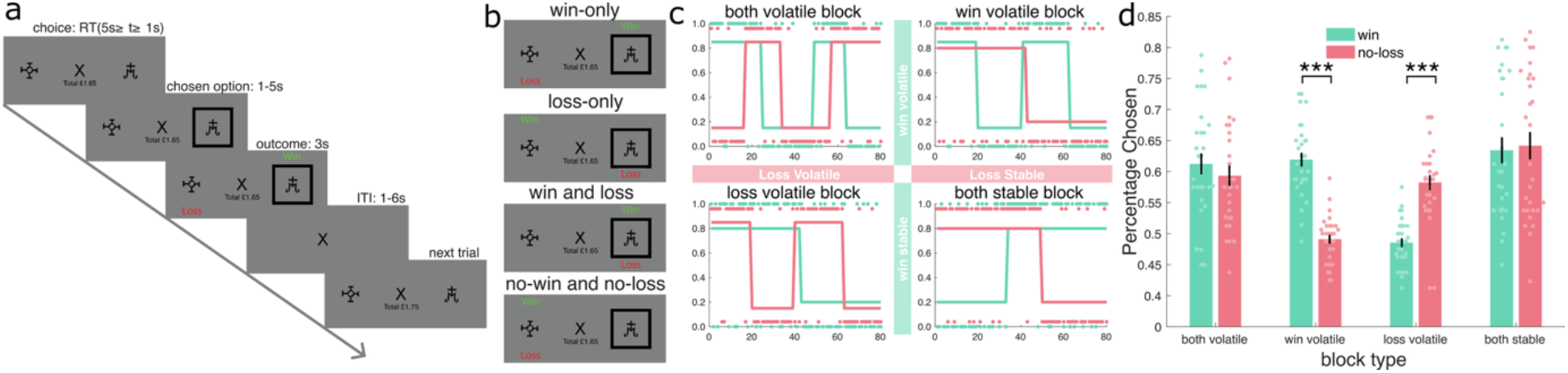
Task design. **a) Timeline of one trial from the information bias learning task**. Participants were presented with two different shapes (shape ‘A’ and ‘B’) and were required to choose one. After a choice, a black frame appeared around the chosen option. After a jittered interval, the outcomes were presented. For each trial, a win and a loss was associated with one of the shapes. If the shape associated with an outcome was chosen, each win would add 15p to the running total, while each loss would deduct 15p from it. The total amount of money won in the task was presented on the lower half of the screen throughout a trial. **b) four possible trial outcomes**. Wins and losses could be associated with the same or with different shapes, which resulted in four potential outcomes for the chosen option: win-only (as shown in the example trial), loss-only, win and loss, no-win and no-loss. **c) Overall task structure**. The task consisted of 4 blocks of 80 trials. One for each block type: both-volatile, win-volatile, loss-volatile, and both-stable. The y-axis represents the probability, p, that an outcome (win in green or loss in red) was be associated with shape ‘A’ (for shape ‘B’ the association is 1-p). The blocks differed in how volatile (changeable) the outcome probabilities were. Block order was counterbalanced across participants. **d) Percentage of win and loss choice accuracies for each block for all the participants**. Each dot represents each participant’s data. Error bars represent ±1 standard error of the mean.

All self-report measures, other than the Temporal Experience of Pleasure Scale (TEPS) (Gard, Gard et al. 2006), were very highly correlated with each other (see Fig. S1), suggesting that the scores were largely reflective of general negativity, rather than specific symptom clusters (Gagne, Zika et al. 2020). Consistent with this, factor analysis of questionnaire scores indicated a dominant single factor that explained variance across all measures (only this single factor exceeded Kaiser’s criterion, i.e. had an eigenvalue of greater than one (see Fig. S2)). To simplify analysis, this first factor score is therefore used as the summary measures of the participants’ overall negativity level, which we referred to as “Negativity scores” in the following text.

### Volatile outcomes dominated participant choice, regardless of valence

In the version of the information bias learning task (Pulcu and Browning 2017) used in the current study (see Methods for more details), participants made a series of forced choices between two options. Following each choice, both a rewarding (win) and a punishing (loss) outcome was presented, associated with one of the two options (see Fig 1). The win and loss outcomes were independent of each other, such that on each trial participants could receive one of four possible outcome combinations: either a win-only, a loss-only, both a win and a loss, or neither a win nor a loss (see Fig 1b). As a result, participants had to separately estimate the win and loss probabilities. The task consisted of four blocks of 80 trials each. In each block the volatility of win and loss was manipulated to be either volatile (probabilities switched between 85 and 15% three times), or stable (probabilities switched from 80 to 20% once; Figure 1c), resulting in four block types: both-volatile, win-volatile (win volatile, loss stable), loss-volatile (loss volatile, win stable) and both-stable blocks (see Methods for more details). Block order was counterbalanced across participants.

We first examined participants’ performance in the task, specifically their choice accuracies for wins and losses (Figure 1d). For balanced blocks (both-volatile and both-stable blocks), participants showed equal performance for wins and losses (valence effect within blocks, both p>.536) and performed above the chance level (one-sample T-test against 50%, all t(28)>5.607, p<.001). In contrast, when there was an imbalance in volatility between outcomes, participant choices were dominated by the more volatile outcome; win accuracy was higher than loss in the win-volatile block (t(28)=9.931, p<.001), and vice versa for loss accuracy in the loss-volatile block (t(28)=-6.850, p<.001) (see Fig. 1d). One possible reason for this is that, to compensate for the difficulty in estimating volatile associations, which frequently change over time, our design followed [Berhens et al 2007] in using slightly more extreme probabilities for the volatile (85%:15%) than stable (80%:20%) outcomes. However, as can be seen in Figure 1d, despite the stable outcomes being associated with participant choice at 80%, the influence of these outcomes on choice accuracy was completely suppressed for both wins in the loss-volatile block (one sample t-test against 50%, t(28)=-2.023, p=.053) and losses in the win-volatile block (t(28)=-1.258, p=.219).

See the supplement (Table S2) for further analysis of the task using an ANCOVA framework. In the following section, we further examine these effects using computational modelling.

### Higher negativity scores are associated with a specific reduction in loss learning rate adaptation to volatility

To characterize how participants adapted their learning to the volatility of the two outcomes, we used a volatility adaptation model in which Rescorla-Wagner rules (Rescorla 1972) were applied to update win and loss values independently. Reflecting the structure of the task, the model (see methods and Table 1 for more detailed description) estimated learning rate parameters for wins and losses when they were stable using two “base” (stable) learning rates for wins in the loss-volatile and both-stable blocks, and two “base” (stable) learning rates for losses in the win-volatile and both-stable blocks. The model accounted for outcome volatility using four learning rate adaptation terms, such that the learning rates for volatile conditions were estimated as the base values plus the relevant volatility adaptation term. This parameterisation allowed the model to directly capture the effects of the volatility manipulation in the volatility adaptation terms and also supported the derivation of win and loss learning rates for each block (see Fig. 2 a).

**Table 1.** Equations for the volatility adaptation model for the fMRI study.

| block | valence | Description | equation |
| --- | --- | --- | --- |
| Both Volatile | win | Win volatile learning rate | $\alpha_{win}(1) = \alpha_{win_{stable2}} + \alpha_{win_{adpt2}}$ |
| | loss | Loss volatile learning rate | $\alpha_{loss}(1) = \alpha_{loss_{stable2}} + \alpha_{loss_{adpt2}}$ |
| Win Volatile | win | Win volatile learning rate | $\alpha_{win}(2) = \alpha_{win_{stable1}} + \alpha_{win_{adpt1}}$ |
| | loss | Loss stable learning rate | $\alpha_{loss}(2) = \alpha_{loss_{stable2}}$ |
| Loss Volatile | win | Win stable learning rate | $\alpha_{win}(3) = \alpha_{win_{stable2}}$ |
| | loss | Loss volatile learning rate | $\alpha_{loss}(3) = \alpha_{loss_{stable1}} + \alpha_{loss_{adpt1}}$ |
| Both Stable | win | Win stable learning rate | $\alpha_{win}(4) = \alpha_{win_{stable1}}$ |
| | loss | Loss stable learning rate | $\alpha_{loss}(4) = \alpha_{loss_{stable1}}$ |

Note: Stable 1 refers to stable parameters when other-volatility was stable; Stable 2 refers to stable parameters when other-volatility was volatile; adpt1 refer to adaptation terms when other-volatility was stable; adpt2 refer to adaptation terms when other-volatility was volatile.

**Fig. 2.**
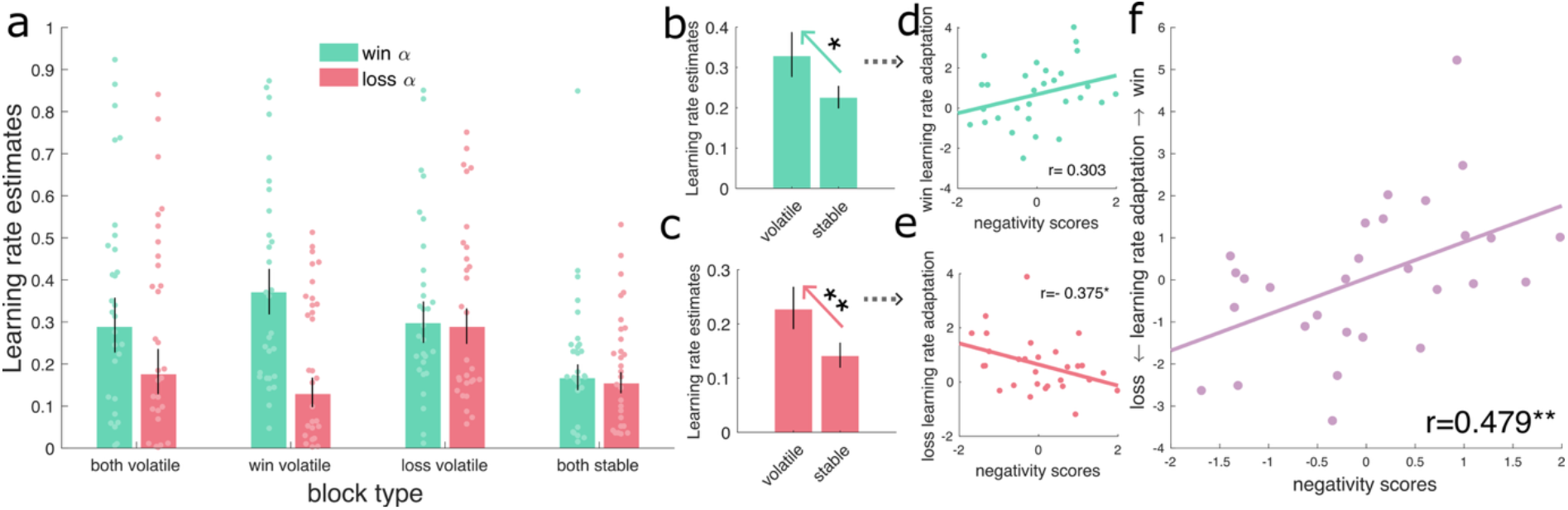
Valence bias in learning rate adaptations correlated with overall negativity scores. a) Win and loss learning rate estimates for each block. b) An illustration of the volatility adaptation effect for wins, average win learning rates for win volatile blocks (i.e. both volatile and win volatile) are displayed besides average win learning rates for win stable blocks (i.e. loss volatile and both stable). c) An illustration of the volatility adaptation effect for losses, average loss learning rates for loss volatile blocks (i.e. both volatile and loss volatile) are displayed besides average loss learning rates for loss stable blocks (i.e. win volatile and both stable) d) a scatter plot showing the non-significant positive correlation between win learning rate volatility adaptation and the overall negativity scores. e) a scatter plot showing the significant negative correlation between loss learning rate volatility adaptation and the overall negativity scores. f) a scatter plot illustrating the relative win vs. loss learning rate volatility adaptation bias positively correlated with the overall negativity scores. Each dot represents a participant. Error bars represent ±1 standard error of the mean. * p<0.05, **p<0.01.

We first examined whether participants increased learning rates (LR) in general for volatile compared to stable environments for both win and loss outcomes. Both win learning rate adaptation (i.e. average of the two win learning rate adaptation terms; t(28)=2.410, p=.023, see Fig. 2 b) and loss learning rate adaption (i.e. average of the two loss learning rate adaptation terms; (t(28)=3.383, p=.002, see Fig. 2 c) were significantly above zero, confirming this basic effect.

We then examined whether LR volatility adaptation significantly differed between wins and losses, that is, whether there was a valence bias in volatility adaptation. A paired t-test showed that the difference between win and loss LR adaptation was not significant (t(28)=.121, p=.905). This suggested that, in general, participants didn’t show a valence bias in LR volatility adaptation. To support this claim, we ran a Bayesian paired samples t-test, which indicated that the data supported the null hypothesis (H0, i.e., no valence bias) being true (BF_10_ = 0.199, error rate 0.032%). These results suggest that overall people can increase their LR to win and loss volatility respectively, and to the same extent, when both outcome valences are presented simultaneously.

We then examined whether individual differences in win and loss LR adaptations or the valence bias (that is win adaptation – loss adaptation) were associated with questionnaire-based negativity scores. We found a positive but not significant trend level correlation between win LR adaptation and negativity scores (r(27)=0.303, p=.110, see Fig. 2 d), with an opposite and significant negative correlation between negativity scores and loss LR adaptation (r(27)=-0.375, p=.045, see Fig. 2 e), suggesting a valence bias. Indeed, we found that people with higher negativity scores showed relatively more LR adaptation to win vs. loss volatility (r(27)=0.479, p=.009, see Fig. 2f). Thus, while there wasn’t a general valence bias in volatility adaptation at the population level; individuals with higher negativity scores demonstrated a reduced sensitivity to the volatility of losses, relative to wins.

For clarity, we focus on presenting the results relevant to our primary hypotheses here; details of the full ANCOVA model estimates are available in the Supplement (Table S3 & Table S4).

### The volatility of reward and loss outcomes are represented by activity in distinct subregions of the ACC

The ACC has been shown to track volatility during learning (Behrens, Woolrich et al. 2007, Silvetti, Seurinck et al. 2013) for reward outcomes. Here we examine how win (reward) and loss (punishment) volatility are represented in the ACC when the task requires tracking of both outcome types. We first identified subregions of the ACC that robustly differentiated between positive (win or no-loss) and negative (no-win or loss) outcomes (a mean contrast between win vs no-win and no-loss vs loss contrasts estimated from the model described in the Methods). A cluster in the pgACC showed greater BOLD responses to positive than negative outcomes (Fig. 3a, also see Fig.S 3), while a second cluster in the dorsal ACC (dACC) showed greater BOLD responses to negative than positive outcomes (Fig. 3b and Fig.S 3).

**Fig. 3.**
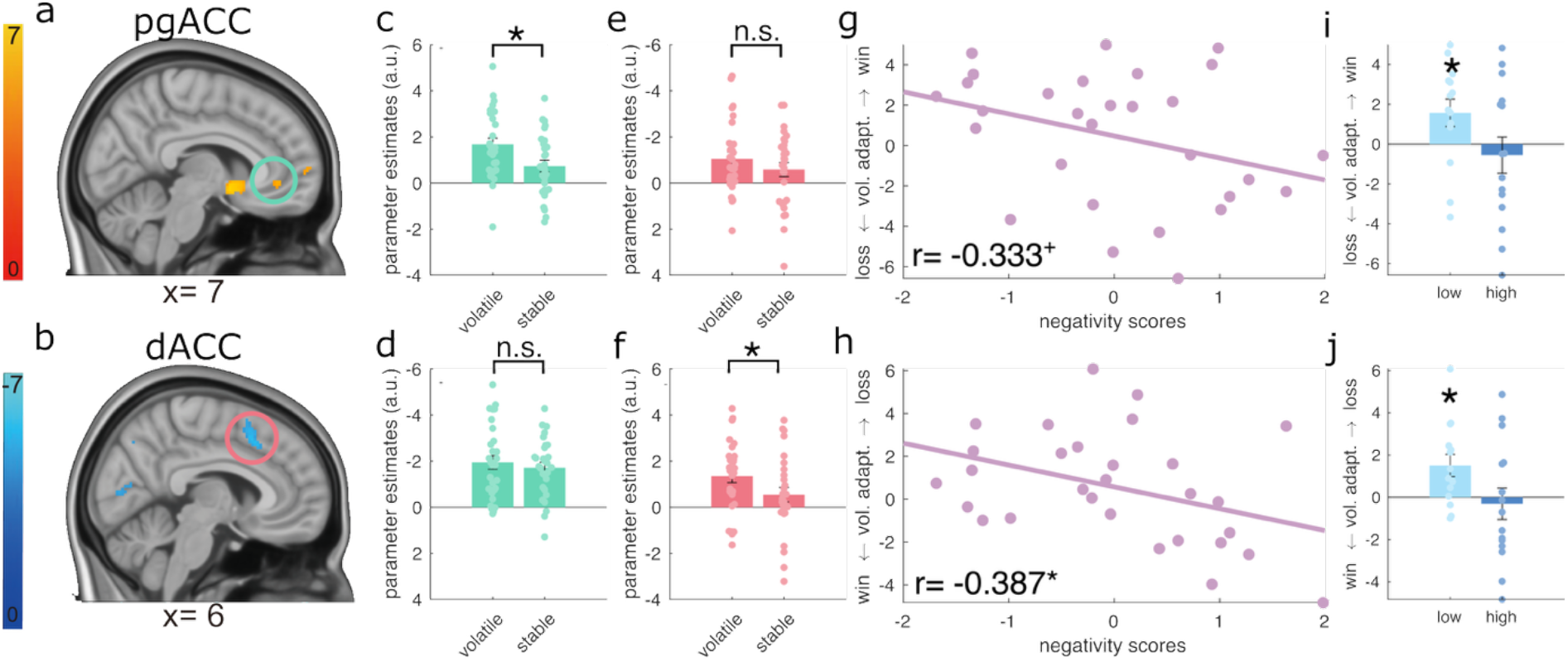
Win and loss volatility signals were distinctly represented in subregions of ACC. a-b) Sagittal views of the perigenual anterior cingulate cortex (pgACC, circled in green) and dorsal (dACC, circled in red). c-d) Extracted BOLD signals for win vs no-win contrast for win volatile and win stable using the pgACC (c) and dACC (d) ROIs. e-f) Extracted BOLD signals for loss vs no-loss contrast for loss volatile and loss stable using the pgACC (e) and dACC (f) ROIs. As can be seen, there is a specific effect of win volatility in the pgACC, and of loss volatility in the dACC. g) the relative adaptation to win vs. loss volatility (win volatile vs. stable – loss volatile vs. stable) in pgACC marginally negatively correlated with the overall negativity scores. i) a bar plot illustrating this effect in pgACC with groups median split by negativity scores. h) the relative adaptation to loss vs. win volatility (loss volatile vs. stable – win volatile vs. stable) in dACC negatively correlated with the overall negativity scores. j) a bar plot illustrating this effect in dACC with groups median split by negativity scores. Error bars represent ±1 standard error of the mean. +p<0.1, *p<0.05, n.s. not significant

To examine whether volatility differentially modulated BOLD responses to win and loss outcomes in these two functionally defined ROIs, i.e. pgACC and dACC (Fig. 3a&b), we entered parameter estimates from the regions for wins (win vs. no-win) and losses (no-loss vs. loss) outcome contrasts in the volatile and stable conditions into a repeated measures ANCOVA (full results see Supplemental Table S5). A significant region*valance*volatility interaction effect was found (F(1,27)=20.566, p<.001, η_p_2=.432), indicating that volatility adaption to rewards and losses differed between the two ACC regions. Post hoc analyses, summarised in Fig. 3, illustrates that this effect was driven by the BOLD response in the pgACC BOLD response to win outcomes being significantly modulated by win volatility (win-volatile vs win-stable, t(28)=2.560, p=.016, Fig. 3c), whereas the dACC response to loss outcomes was significantly influenced by loss volatility (loss-volatile vs loss-stable, t(28)=2.286, p=0.030, Fig. 3f). In other words, the volatility of rewards and losses were tracked by anatomically distinct regions, with activity in the dACC tracking loss volatility and the pgACC tracking win volatility.

### Negativity scores are associated with reduced anatomical specificity in the representation of volatility

The anatomical specificity of volatility adaptation was modulated by participants’ negativity scores (region*valence* volatility*negativity scores: F(1,27)=6.985, p=.014, η_p_2=.206). As can be seen in figure 3g-j, whereas participants with low negativity scores maintained a high degree of anatomical specificity, with the pgACC tracking win rather than loss volatility (Fig. 3i), and the dACC tracking loss rather than win volatility (Fig. 3j), as negativity scores increased, the specificity of the dACC for loss volatility was reduced (r(27)= 0.387, p=.038, Fig. 3h), as was the specificity of the pgACC for win volatility, albeit at a trend level (r(27)= -0.333, p= .077, Fig. 3g).

In the previous results, we described that negativity scores significantly correlated with the relative adaptation of both participant behaviour (learning rates, Fig. 2f) and BOLD signal to the volatility of wins vs. losses in the dACC (Fig. 3h). We therefore examined how the behavioural and neuroimaging volatility adaptation measures might be linked. We found that the specificity of the BOLD responses in the dACC for loss volatility was positively correlated with relative loss vs. win learning rate volatility adaptation (r(27)= 0.413, p=.026, Fig. 4a). There was no correlation with BOLD responses in the pgACC (p=.866). In other words, the more an individual’s dACC specifically tracked loss, relative to win, volatility, the more that person adapted their learning rates to changes in loss, relative to win, volatility. Finally, we tested a mediation model (Fig. 4 b) in which the neuroimaging effects, drive a differential learning rate response, which then alters negativity scores. Specifically, we tested whether loss vs. win learning rate adaptation mediates the relationship between the dACC BOLD response and negativity scores. The causal mediation analysis revealed a significant average mediation effect (ACME=-0.060, CI=[-0.135 -0.002], p=.042), suggesting that behavioural learning rate adaptations mediated the relationship between dACC and negativity scores. A similar analysis of the mediating effect of learning rate adaptation on the relationship between pgACC and negativity was not significant (ACME= -0.005, CI=[-0.057 0.063], p=.885).

**Fig. 4.**
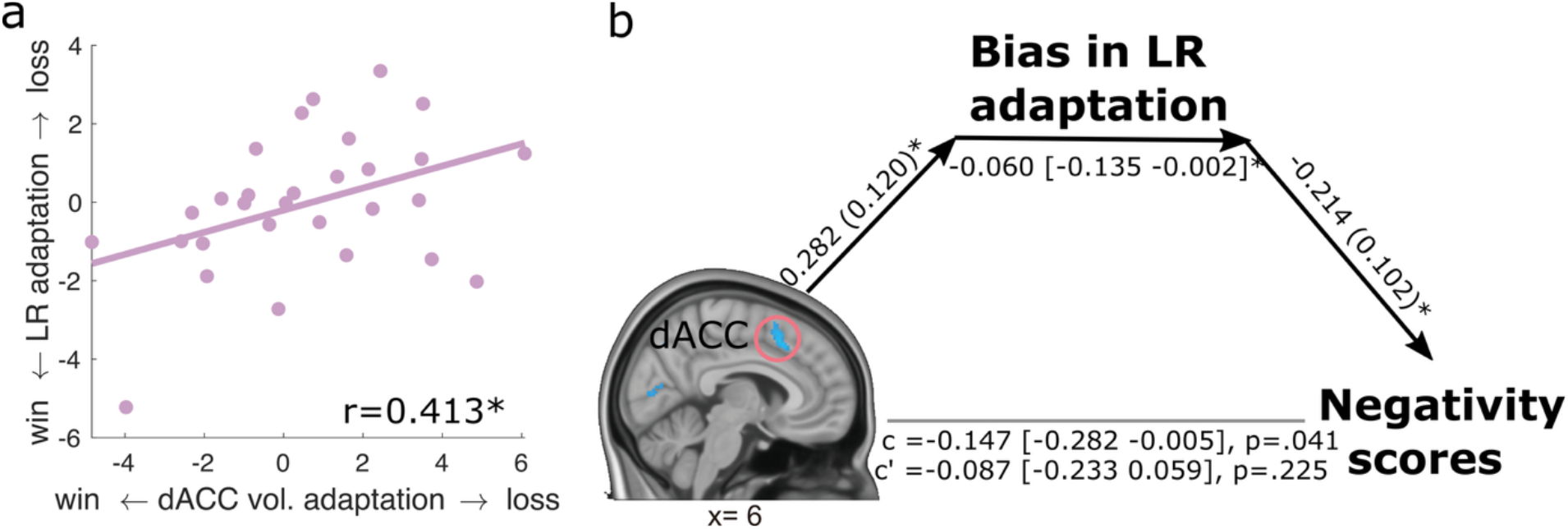
a) a scatter plot showing an association between the relative loss vs. win volatility adaptation bias in the dACC and relative loss vs. win volatility adaptation bias in learning rates (LR). b) A mediation analysis suggests that the loss vs win relative volatility adaptation in dACC modulated overall negativity scores via change of valence bias in learning rate adaptations. As illustrated, the direct path c’ effect is no longer significant. * 0.01<p<0.05

## Discussion

People adapt to environmental volatility by increasing learning rates when outcomes are volatile compared to when they are stable. In this study, we showed that people can independently adjust learning rates for reward (win) and punishment (loss) volatility when both sources of information are present concurrently (Fig. 2). Using fMRI, we demonstrated that win and loss volatility modulate distinct subregions of the ACC: pgACC and dACC (Fig. 3), respectively. However, individuals with higher overall negativity scores showed a relative bias in learning rate adaptation. They adapted learning more to wins compared to losses, driven by a reduced learning rate adaptation to losses. This behavioural bias was associated with less distinct neural encoding of win and loss volatility in the pgACC and dACC. The association between reduced loss volatility signalling in the dACC and negativity scores was found to be, mediated via the valence bias in learning rate adaptation (Fig. 4). Together, our findings describe a neural mechanism that allows the independent tracking of reward and punishment volatility. This suggests that the reduced volatility adjustment in individuals with higher levels of anxiety (Browning, Behrens et al. 2015, Pulcu and Browning 2017, Gagne, Zika et al. 2020) and depression may arise because these individuals struggle to maintain distinct representations of the volatility of valenced outcomes.

Our finding that people can independently increase learning rates for volatile wins and losses is consistent with prior work (Pulcu and Browning 2017). However, a limitation of this prior work was that the stable schedule used a fixed 50% outcome probability. In addition to producing low unexpected uncertainty (i.e. low volatility), this resulted in a high expected uncertainty (Yu and Dayan 2005). That is, these stable outcomes were uninformative about which choice was best. Here, we designed the stable condition with a single contingency shift per block, to reduce difference with the expected uncertainty of the volatile condition (Fig 1c). This allowed manipulation of unexpected uncertainty (volatility) effects without confounding effects of expected uncertainty. Even with this more learnable stable schedule, participants’ choices and learning rates were dominated by the more volatile source of information, confirming that volatility drives learning adjustments.

We found that people with higher negativity scores showed a relative bias in volatility adaptation, such that they were more responsive to changes in win than loss volatility. On a more granular level, participants with high levels of negativity showed significantly reduced volatility adaptation to losses and a trend-level increase in win volatility adaptation. This is consistent with earlier studies linking anxiety to impaired volatility adaptation for punishments like electric shocks (Browning, Behrens et al. 2015) and monetary losses (Pulcu and Browning 2017). The same impairment in learning adaptation to punishments (electric shocks) was also shown in patients with generalized anxiety disorder (GAD) and major depressive disorder (MDD) (Gagne, Zika et al. 2020). However, they also reported reduced reward volatility adaptation in a separate reward-only task, whereas we observed a trend in the opposite direction in our dual-information task. This discrepancy may arise because, in our task, participants had to integrate and trade off both reward and punishment signals on every trial. The need to balance concurrent valenced information may alter how volatility adaptations manifest, particularly in populations with affective symptoms. This highlights the importance of studying learning in ecologically richer contexts where multiple affective signals co-occur.

Our fMRI results indicated that volatility modulates outcome-related BOLD responses in the ACC, consistent with prior work in humans (Behrens, Woolrich et al. 2007, Silvetti, Seurinck et al. 2013) and non-human primates (Massi, Donahue et al. 2018). However, by presenting participants with cues associated with both reward and loss outcomes on each trial, and by independently manipulating the volatility of these outcomes, we were able to directly probe neural systems displaying valence-specific volatility signals. We found that reward and punishment volatility modulated activity in anatomically distinct ACC subregions. While previous studies have reported valence-based dissociations of the prefrontal cortex during probabilistic learning (Monosov and Hikosaka 2012, Monosov 2017, Gueguen, Lopez-Persem et al. 2021), our study demonstrates a valence-specific volatility effect within the ACC. This distinct neural representation may be crucial for flexibly and independently adapting learning to rewards and punishments in dynamic environments.

Importantly, we found that valence-specific volatility encoding in the ACC subregions was less distinct in individuals with higher negativity scores. Further analyses suggested that the reduced specificity of the dACC for negative volatility may impact individuals’ anxiety and depression via a mediating effect on affecting learning rates. Altered ACC structure and function have long been implicated in mood and anxiety disorders (Myers, Simon et al. 2025). A recent review highlighted dACC as an overlapping region in pathological anxiety and processing unpredictable threats (Chavanne Alice and Robinson Oliver 2021). Dorsal ACC is involved in conflict resolution with interfering negative valence (angry and fearful faces), whose activity is linked to rumination symptoms in depressed and anxiety patients (Sheena, Jimmy et al. 2021). Notably, using a cost-benefit trade-off task similar to ours, in which macaques made a decision about whether to approach or avoid an offer associated with both rewards and punishments (Amemori and Graybiel 2012), a region of the ACC with predominantly negative valenced neurons has been described. Critically, this work demonstrated that stimulating this area biased the animals to more avoidance choices, akin to excessive avoidance - a core feature of anxiety (Ball and Gunaydin 2022), while anxiolytic drug treatment blocked this effect. Taken together, these results suggest negative valenced ACC regions might be particularly important for modulating anxiety-like behaviours which could further ameliorate anxiety and depression symptoms.

A number of limitations to the current study should be acknowledged. While our findings suggest that a potential mechanism for the previously described reduced adaptation to negative volatility in anxiety and depression (Browning, Behrens et al. 2015, Pulcu and Browning 2017, Gagne, Zika et al. 2020) might be a difficulty in maintaining distinct estimates of the statistics of valenced outcomes, an alternative possibility is that this difficulty is one facet of a more general deficit in accurately representing distinct estimates of environmental processes, be they valenced or otherwise. Testing this will require designs in which the volatility of other forms of association are manipulated. Secondly, while this study purposively recruited participants to have a range of different depressive scores, this was not a clinical sample and therefore the relevance of these findings to patients with diagnoses of anxiety and depression is uncertain.

In summary, we have demonstrated that people can independently adapt learning to concurrent reward and punishment volatility, likely supported by distinct neural representations of win and loss volatility in ACC subregions. Impairments in this mechanism—in terms of both behavioural adaptation to loss volatility and a reduced neural separation of the volatility signals associated with positive and negative outcomes—are associated with higher anxiety and depression symptoms. This work underscores the importance of studying learning in multi-valenced environments and suggests that difficulties in maintaining distinct estimates of environmental processes may be a key mechanism in affective disorders.

## Methods

### Participants and experimental protocol

#### Procedures

Participants were screened using the Quick Inventory of Depressive Symptoms (QIDS) online, which measures symptoms of depression (Rush, Trivedi et al. 2003). We then invited participants to the fMRI study based on their QIDS scores, ensuring a roughly equal number of participants with scores 1-5 (no depression), 6-10 (mild depression), and 11-15 (moderate depression). Participants were also selected to ensure that age and gender did not confound the QIDS score (i.e., selection ensured a lack of significant relationship between QIDS scores and either age or gender).

During the fMRI session, participants completed four blocks of the information bias learning task within the MRI scanner. On the scan day, participants were also asked to answer the QIDS again as well as three other questionnaires; (1) the Ruminative Response Scale, which measures rumination in response to depressive mood (RRS) (Nolen-Hoeksema and Morrow 1991), (2) the Spielberger State-Trait Anxiety Inventory (STAI), which measures state and trait anxiety (Spielberger 1983), (3) The Temporal Experience of Pleasure Scale (TEPS), which measures individual trait dispositions in anticipatory and consummatory experiences of pleasure (Gard, Gard et al. 2006). These questionnaires measure different aspects of depressive experience but are generally highly correlated with one another. To account for this, we ran a factor analysis to derive an overall quantification of negative symptoms, using the four total questionnaire scores taken on the scan day.

This study received ethical approval from the Medical Sciences Interdivisional Research Ethics Committee of the University of Oxford’s Central University Research Ethics Committee with the reference R56549/RE002, in accordance with the Declaration of Helsinki. All participants provided written informed consent during their visit.

#### The information bias learning task

The learning task used in this study was adapted from the same learning task described in a previous study in our lab (Pulcu and Browning 2017). In this task, participants were asked to repeatedly choose between two abstract shapes within a block. One of the shapes was associated with a probability of getting a win (P_win_), and an independent probability of getting a loss (P_loss_). The other shape would be associated with a win probability of 1-P_win_, and with a loss probability of 1-P_loss_. Each win and loss had a magnitude of 15p. Therefore, the chosen shape in a given trial might end up with a win and no loss (gain 15p), a loss and no win (lose 15p), a win and a loss (no monetary change), and lastly, no win and no loss (also no monetary change).

On each trial, participants were first presented with the two shapes, during which they were asked to choose between the two shapes. Following this choice, a black frame would appear around the shape that was chosen. Participants’ choices were self-paced. The choice phase was limited to between 1s and 5s. If a participant responded before 1 second, the black frame appeared 1 second after the onset of the choice phase. If a participant did not respond within 5 seconds, a random choice was made for the participant for this trial. The chosen option was marked with a black frame for a jittered interval of 1-5 (mean 3) seconds. The win and loss outcomes were shown at the same time for each trial for a fixed duration of 3 seconds. Finally, the trials were separated with a jittered interval of 1–6 (mean 3.5) seconds. The total monetary winning was shown under the fixation cross and updated at the end of each trial.

Probabilities of wins and losses change in different volatility levels during a block. In a “volatile” schedule, the outcomes probabilities for a given shape switched frequently between 15 and 85% during a block. In a “stable” schedule, the outcomes probabilities for a given shape switched only once between 20 and 80% during a block. Note that the critical change we made to this version of the task compared to the one described by Pulcu et., al (Pulcu and Browning 2017) is that we made the “stable” schedule as a single switch between high and low probabilities, which was fixed at 50% in the original paper. Therefore, the stable schedule in this version of the task was still learnable but relatively stable in terms of contingency changeability. This reduced the difference in expected uncertainty between the volatile and stable schedules, allowed us to add a both-stable block, which was not in the previous version, and enabled us to test the valence*self-volatility*other-volatility effect in our analysis.

Critically, to test how the volatility of the two independent information (win and loss) affects the learning of each of the information per se and how they interact, we arranged the four blocks in a valence (win or loss)*volatility (volatile and stable) design. More specifically, the four blocks are 1) the both-volatile block, where both win and loss are volatile, 2) the win-volatile block, where win is volatile and loss is stable, 3) the loss-volatile block, where loss is volatile and win is stable, 4) the both-stable block, where both win and loss are stable. Participants need to track both the win and loss probabilities to maximize their monetary payoff. Participants were reimbursed with a base amount plus the money they earned from the task. The participants were also informed about the bonus and that the win and loss contingencies were independent and might change during a block.

The block orders were balanced across participants. To avoid any unbalanced volatility manipulation between win and loss schedules, win schedules for the half of the participants were used as loss schedules for the other half of the participants, and vice versa for the loss schedules. We also reverse the win and loss schedules for the semi-volatile block within a participant, i.e., the win schedule in the win volatile block were used as loss schedule in the loss volatile block, vice versa for the loss schedule.

### Computational models

#### The volatility adaptation model

The expected values for win and loss were updated trial by trial with block-wise win and loss learning rate respectively using a Rescorla-Wagner learning rule (Rescorla 1972) (see Equation 1, Equation 2). The Q value was set to 0.5 for both win and loss probabilities at the beginning of each block. These expected values were then transformed into a single choice probability using a softmax function (see Equation 3).

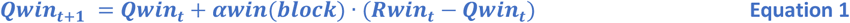

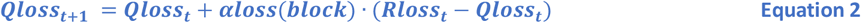

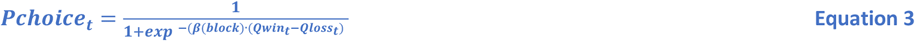

*α*_stable_ denotes the base learning rates. *α*_adpt_ denotes the learning rate differences or adaptations from stable to volatile environments. To account for the background environment (how volatile is the other available information), we fitted separate sets of stable and volatile adaptation learning rates for blocks in which the other outcome was volatile (set 2) vs. stable (set 1) (see Table 1), and different inverse temperatures for each block. The first 5 trials of each block were not used in model fitting.

#### The beta adaptation model

The betas volatility adaptation model assigned different betas for each block for wins and losses respectively but with the same learning rates for wins and losses for each block (see Table 2). The model assumes that volatility only affects how participants use win and loss information to make decisions but not learning rate per se.

**Table 2.** Equations for the beta volatility adaptation model for the clinical study.

| block | valence | Description | equation |
| --- | --- | --- | --- |
| Both Volatile | win | Win volatile beta | $\beta_{win}(1) = \beta_{win_{stable}} + \beta_{win_{adpt}}$ |
| | loss | Loss volatile beta | $\beta_{loss}(1) = \beta_{loss_{stable}} + \beta_{loss_{adpt}}$ |
| Win Volatile | win | Win volatile beta | $\beta_{win}(2) = \beta_{win_{stable}} + \beta_{win_{adpt}}$ |
| | loss | Loss stable beta | $\beta_{loss}(2) = \beta_{loss_{stable}}$ |
| Loss Volatile | win | Win stable beta | $\beta_{win}(3) = \beta_{win_{stable}}$ |
| | loss | Loss volatile beta | $\beta_{loss}(3) = \beta_{loss_{stable}} + \beta_{loss_{adpt}}$ |

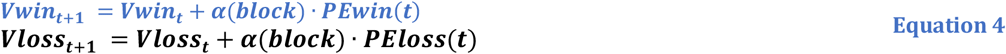

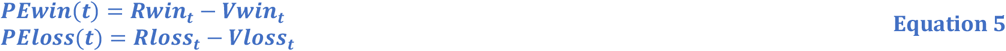

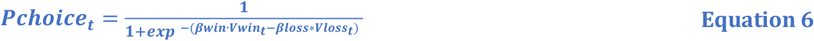

#### The fixed learning model

The fixed learning model used only one learning rate and one beta for all the blocks. The model assumes that neither learning rates nor betas were affected by volatility manipulations.

#### Model fitting procedure

All models were fitted using Stan (RStan v 2.21.1, R v 3.6.0), a software package that uses Markov chain Monte Carlo method to sample from posterior distributions of parameters (Carpenter, Gelman et al. 2017). 4 parallel Monte Carlo chains were run each with 5000 iterations (5000 warm-ups), using adapt_delta of 0.99 and maximum treedepth of 10. Learning rate parameters were sampled from a zero-mean normal distribution with a standard deviation of 2, and then transformed using logistic approximation (Bowling, Khasawneh et al. 2009). Beta parameters were sampled from a zero mean normal distribution with standard deviation 1, and then transformed using the exp function. Convergence was inspected using shinyStan (v 2.5.0). No convergence problems were detected. We adopted a stringent cut-off of R-hat< 1.1 as an indicator for sufficient convergence (Brooks and Gelman 1998), and the minimum number of effective samples >50. The models were fitted to each participant’s data without a group hierarchical setting to optimise for individual differences analyses.

We then performed model comparisons using a leave-one-out cross-validation procedure using the LOO package for R (Vehtari, Gelman et al. 2017). An expected log pointwise predictive density (ELPD) and a standard error for each ELPD estimate were generated for each model as a measure of predictive accuracy.

### fMRI data analysis

#### fMRI data acquisition and pre-processing

All image data was acquired using a Siemens Prisma 3 T scanner (the Oxford Centre for Human Brain Activity, University of Oxford, Oxford, UK) with a 32-channel head coil. The task-based fMRI images were acquired using a multiband-echo planar imaging (mb-EPI) sequence, with a multiband factor of 6, repetition time (TR) = 800 ms, echo time (TE) = 30 ms, 60 slices (no gap), a slice angle of 30°, interleaved acquisition, voxel size = 2.4 × 2.4 × 2.4 mm, flip angle = 52°, and field of view (FOV) = 216 mm, with local z-shimming). Field maps were obtained using a dual echo 2D gradient-echo sequence with TR = 590 ms, TE = 7.38 ms, and 4.92ms, and flip angle = 46°. The high-resolution T1-weighted anatomical MRI images were acquired using a 3D MPRAGE sequence with TR = 1900 ms, TE = 3.97 ms, 192 transversal slices, voxel size = 1 ×1 × 1 mm, and FOV = 192 mm.

The fMRI data were pre-processed using FEAT (FMRI Expert Analysis Tool) part of FSL (FMRIB Software Library V6 (Analysis Group, FMRIB, Oxford, UK; http://www.fmrib.ox.ac.uk/fsl) (Jenkinson, Beckmann et al. 2012). The procedures included (1) motion correction to a high-resolution functional image acquired at the beginning of each block using MCFLIRT (Jenkinson, Bannister et al. 2002), (2) spatial correction for magnetic field inhomogeneities estimated by the B0 fieldmap (Jenkinson 2004), (3) brain extraction using BET (Smith 2002), (4) spatial smoothing using a Gaussian kernel (FWHM 5 mm), (5) high-pass temporal filtered (90 s cut-off). Then functional images were registered to both the high-resolution T1w structural scan and standard space using FLIRT.

#### First-level and group-level analysis

The first-level and group-level GLM whole-brain and ROI-based analysis were conducted using FEAT (note, functionally defined ROIs were used, and are described below). We constructed the event-based general linear model (GLM) with the following parametric regressors at choice and outcome onset for each block. At choice onset: (1) the response (coded as 1 and -1, for the right or left response, respectively), (2) choice reaction time, and (3) the trial-by-trial decision (coded as 1 if subject switched away from the previous choice or -1 if she stayed with the same option), (4) choice mean: value set at 1. At outcome onset: (5) win outcome (coded as 1 and -1, for the win received or win not-received, respectively.), (6) loss outcome (coded as 1 and -1, for the loss received or loss not-received, respectively.), (4) outcome mean: value set at 1. Note that the trials to which participants failed to respond in time were coded using the mean of each regressor. (Note that here we used actual outcome results but not PEs to avoid the confound of model estimates. However, we have also run a similar analysis with PEs instead of outcomes. The results were similar.) The regressors were then demeaned before entering into Feat. Standard motion parameters estimated using MCFLIRT motion correction and physiological noise regressors estimated using Physiological Noise Modelling (PNM) tool (Brooks, Beckmann et al. 2008) were also added in the GLM to regress out the effects of head motion and physiological noise respectively.

We first identified regions that might represent outcome signals in the brain. We estimated the mean of the parameter estimates for the contrast for positive outcomes (i.e. receiving a win and not receiving a loss) vs. negative outcomes (i.e. not receiving a win and receiving a loss) across all the 4 blocks at the individual level and then brought them to the whole-brain group-level analysis (one sample t-test). The images were cluster-thresholded at z > 4.6 to identify robustly specific ROIs, and clusters shown survived a statistical test for extent (p < 0.05, fully corrected for multiple comparisons).

To examine whether these outcome signals in the brain were modulated by valence, self-volatility, or other volatility and how overall negativity modulated the effects, we extracted the parameter estimates for the clusters identified by the positive vs. negative outcomes contrast described above, for each block, for win and loss outcomes and for each participant respectively. An ANCOVA, with valence, self-volatility and other-volatility as repeated measures and the overall negativity scores as a covariate, were performed for each region. We followed up the significant effects with paired t-tests. For any significant effect of the negativity scores, we perform Pearson correlations to further examine the covariations.

#### ROI mediation analysis

A causal mediation analysis was performed using the mediation package in R (Tingley, Yamamoto et al. 2014), with non-parametric bootstrapping (n=5000) was performed to further examine the relationships amongst the neural signals in the dACC, learning rate estimates and the negativity traits. In the analysis, the loss vs. win learning rate volatility adaptation bias was set as the mediation factor, and the relative adaption to loss vs. win volatility signals in the dACC and the overall negativity scores as independent and dependent variables, respectively. The same analysis was repeated for win vs loss volatility signals in rACC.

### Statistical analysis

The model-free behavioural data were analysed using MATLAB (version R2018b; MathWorks, MA, USA) and SPSS (version 26.0; IBM, NY, USA). The percentages of positive outcomes (wins or no-loss) chosen were calculated for each block. As proportion data, such as percentages, violate the assumptions of equality of variance required by most parametric tests, the percentage data were arcsine square-root transformed before being submitted to statistical tests. We first examined whether the performance for each block was significantly above chance level (i.e., 50%) using one-sample t-tests. Valence differences (win vs loss) within the block were examined using paired t-tests.

To test the effect of self-volatility on learning rate adaptation (i.e. learning rates should be higher when an outcome is volatile), we ran one-sample t-tests to check whether the learning rate adaptation terms were significantly higher than zero. Bayesian one-sample t-tests were performed using JASP software (Version 0.19.3) to quantify evidence for/against null hypotheses.

Extracted parameter estimates of the win - no win and loss – no loss contrasts from the fMRI analysis were entered into a repeated measures ANCOVA, with within subject factors of: region (pgACC vs dACC), valence (win vs loss) and volatility (volatile vs stable) and between subject term for negativity score (covariate variable).

## Acknowledgements

This study was funded by an MRC clinician scientist fellowship to M.B. (MR/N008103/1). M.B. was also supported by the NIHR Mental Health Translational Research Collaboration and the Oxford Health Biomedical Research Collaboration. The views expressed are those of the authors and not necessarily those of the National Health Service, the NIHR, or the Department of Health and Social Care in the UK. E.P. was funded by Oxford NIHR Oxford Biomedical Research Center. L.T.H. was supported by Henry Dale fellowship 208789/Z/17/Z) and a sLoLa grant from the Biotechnology and Biological Sciences Research Council (BB/W003392/1). W.L. was also supported by a grant from the Lundbeck Foundation (R483-2024-1606).

## Author contributions

M.B., E.P., and W.L. designed research; W.L. and E.P. performed research; W.L. analyzed data; and W.L., M.B., E.P., and L.T.H. wrote the paper.

## Competing interests

M.B. has received consulting fees from Alto Neuroscience, Engrail Neuroscience, Empyrean Neuroscience, and Janssen. M.B. has acted as a previously owned shares in P1vital Ltd. M.B. and W.L. have received travel expenses from Lundbeck for attending conferences. W.L.’s current position is supported by a grant from Lundbeck Foundation. E.P. and L.T.H do not have any competing interests to disclose.

## References

Amemori, K.-i. and A. M. Graybiel (2012). “Localized microstimulation of primate pregenual cingulate cortex induces negative decision-making.” Nature Neuroscience 15(5): 776–785.

Ball, T. M. and L. A. Gunaydin (2022). “Measuring maladaptive avoidance: from animal models to clinical anxiety.” Neuropsychopharmacology 47(5): 978–986.

Behrens, T. E., M. W. Woolrich, M. E. Walton and M. F. Rushworth (2007). “Learning the value of information in an uncertain world.” Nature neuroscience 10(9): 1214–1221.

Behrens, T. E. J., L. T. Hunt, M. W. Woolrich and M. F. S. Rushworth (2008). “Associative learning of social value.” Nature 456(7219): 245–249.

Blain, B. and R. B. Rutledge (2020). “Momentary subjective well-being depends on learning and not reward.” eLife 9: e57977.

Bowling, S. R., M. T. Khasawneh, S. Kaewkuekool and B. R. Cho (2009). “A logistic approximation to the cumulative normal distribution.” Journal of industrial engineering and management 2(1): 114–127.

Brooks, J. C., C. F. Beckmann, K. L. Miller, R. G. Wise, C. A. Porro, I. Tracey and M. Jenkinson (2008). “Physiological noise modelling for spinal functional magnetic resonance imaging studies.” Neuroimage 39(2): 680–692.

Brooks, S. P. and A. Gelman (1998). “General methods for monitoring convergence of iterative simulations.” Journal of computational and graphical statistics 7(4): 434–455.

Browning, M., T. E. Behrens, G. Jocham, J. X. O’reilly and S. J. Bishop (2015). “Anxious individuals have difficulty learning the causal statistics of aversive environments.” Nature neuroscience 18(4): 590–596.

Carpenter, B., A. Gelman, M. D. Hoffman, D. Lee, B. Goodrich, M. Betancourt, M. Brubaker, J. Guo, P. Li and A. Riddell (2017). “Stan: A probabilistic programming language.” Journal of statistical software 76(1).

Chavanne Alice, V. and J. Robinson Oliver (2021). “The Overlapping Neurobiology of Induced and Pathological Anxiety: A Meta-Analysis of Functional Neural Activation.” American Journal of Psychiatry 178: 156–164.

De Berker, A. O., R. B. Rutledge, C. Mathys, L. Marshall, G. F. Cross, R. J. Dolan and S. Bestmann (2016). “Computations of uncertainty mediate acute stress responses in humans.” Nature communications 7: 10996.

Drevets, W. C., J. Savitz and M. Trimble (2008). “The subgenual anterior cingulate cortex in mood disorders.” CNS spectrums 13(8): 663.

Fujiwara, J., P. N. Tobler, M. Taira, T. Iijima and K.-I. Tsutsui (2009). “Segregated and integrated coding of reward and punishment in the cingulate cortex.” Journal of neurophysiology 101(6): 3284–3293.

Gagne, C., O. Zika, P. Dayan and S. J. Bishop (2020). “Impaired adaptation of learning to contingency volatility in internalizing psychopathology.” eLife 9: e61387.

Gard, D. E., M. G. Gard, A. M. Kring and O. P. John (2006). “Anticipatory and consummatory components of the experience of pleasure: A scale development study.” Journal of Research in Personality 40(6): 1086–1102.

Gueguen, M. C. M., A. Lopez-Persem, P. Billeke, J.-P. Lachaux, S. Rheims, P. Kahane, L. Minotti, O. David, M. Pessiglione and J. Bastin (2021). “Anatomical dissociation of intracerebral signals for reward and punishment prediction errors in humans.” Nature Communications 12(1): 3344.

Holroyd, C. B. and N. Yeung (2012). “Motivation of extended behaviors by anterior cingulate cortex.” Trends in Cognitive Sciences 16(2): 122–128.

Jenkinson, M. (2004). “Improving the registration of B0-disorted Epi images using calculated cost function weights: we 202.” Neuroimage 22.

Jenkinson, M., P. Bannister, M. Brady and S. Smith (2002). “Improved optimization for the robust and accurate linear registration and motion correction of brain images.” Neuroimage 17(2): 825–841.

Jenkinson, M., C. F. Beckmann, T. E. Behrens, M. W. Woolrich and S. M. Smith (2012). “Fsl.” Neuroimage 62(2): 782–790.

Klein-Flügge, M. C., A. Bongioanni and M. F. S. Rushworth (2022). “Medial and orbital frontal cortex in decision-making and flexible behavior.” Neuron 110(17): 2743–2770.

Koss, K. J., S. Kronaizl, R. Brown and J. Brooks-Gunn (2025). “Childhood Environmental Unpredictability and Adolescent Mental Health and Behavioral Problems.” Child Development.

Marschark, E. D. and R. Baenninger (2002). “Modification of instinctive herding dog behavior using reinforcement and punishment.” Anthrozoös 15(1): 51–68.

Massi, B., C. H. Donahue and D. Lee (2018). “Volatility Facilitates Value Updating in the Prefrontal Cortex.” Neuron 99(3): 598–608.e594.

Monosov, I. E. (2017). “Anterior cingulate is a source of valence-specific information about value and uncertainty.” Nature Communications 8(1): 134.

Monosov, I. E. and O. Hikosaka (2012). “Regionally Distinct Processing of Rewards and Punishments by the Primate Ventromedial Prefrontal Cortex.” The Journal of Neuroscience 32(30): 10318.

Myers, D. C., J. Simon, J. Oh, K. E. Kabotyanski, S. Fujimoto, D. J. Oathes, P. H. Rudebeck, K.-i. Amemori, S. A. Sheth and J. L. Fudge (2025). “Circuit-Based Approaches to Understanding the Anterior Cingulate Cortex (ACC).” The Journal of Neuroscience 45(46): e1311252025.

Nolen-Hoeksema, S. and J. Morrow (1991). “A prospective study of depression and posttraumatic stress symptoms after a natural disaster: the 1989 Loma Prieta Earthquake.” Journal of personality and social psychology 61(1): 115.

Price, J. L. and W. C. Drevets (2012). “Neural circuits underlying the pathophysiology of mood disorders.” Trends in cognitive sciences 16(1): 61–71.

Pulcu, E. and M. Browning (2017). “Affective bias as a rational response to the statistics of rewards and punishments.” Elife 6: e27879.

Rescorla, R. A. (1972). “A theory of Pavlovian conditioning: Variations in the effectiveness of reinforcement and nonreinforcement.” Current research and theory: 64–99.

Ross, L. T., B. Heming and A. Lane (2023). “Family unpredictability and sense of coherence: Relationships with anxiety and depression in two samples.” Psychological Reports 126(4): 1701–1724.

Ross, L. T., C. O. Hood and S. D. Short (2016). “Unpredictability and symptoms of depression and anxiety.” Journal of Social and Clinical Psychology 35(5): 371–385.

Rush, A. J., M. H. Trivedi, H. M. Ibrahim, T. J. Carmody, B. Arnow, D. N. Klein, J. C. Markowitz, P. T. Ninan, S. Kornstein and R. Manber (2003). “The 16-Item Quick Inventory of Depressive Symptomatology (QIDS), clinician rating (QIDS-C), and self-report (QIDS-SR): a psychometric evaluation in patients with chronic major depression.” Biological psychiatry 54(5): 573–583.

Sheena, M. K., J. Jimmy, K. L. Burkhouse and H. Klumpp (2021). “Anterior cingulate cortex activity during attentional control corresponds with rumination in depression and social anxiety.” Psychiatry Research: Neuroimaging 317: 111385.

Sidin, S. A. (2021). The application of reward and punishment in teaching adolescents. Ninth International Conference on Language and Arts (ICLA 2020), Atlantis Press.

Silvetti, M., R. Seurinck and T. Verguts (2013). “Value and prediction error estimation account for volatility effects in ACC: A model-based fMRI study.” Cortex 49(6): 1627–1635.

Smith, S. M. (2002). “Fast robust automated brain extraction.” Human brain mapping 17(3): 143–155.

Soltani, A. and A. Izquierdo (2019). “Adaptive learning under expected and unexpected uncertainty.” Nature Reviews Neuroscience 20(10): 635–644.

Spielberger, C. D. (1983). “State-trait anxiety inventory for adults.”

Tingley, D., T. Yamamoto, K. Hirose, L. Keele and K. Imai (2014). “mediation: R Package for Causal Mediation Analysis.” Journal of Statistical Software 59(5): 1–38.

Vehtari, A., A. Gelman and J. Gabry (2017). “Practical Bayesian model evaluation using leave-one-out cross-validation and WAIC.” Statistics and Computing 27(5): 1413–1432.

Yu, A. J. and P. Dayan (2005). “Uncertainty, Neuromodulation, and Attention.” Neuron 46(4): 681–692.

