## supplementary info for "Valence-specific representation of uncertainty in the anterior cingulate is impaired in those with affective symptoms"

Indistinct representation of reward and punishment uncertainty in those with affective symptoms

### Supplement Methods

### Supplement results

**Table. S 1| Demographic details of the participants**

| variables | Mean | SD |
|---|---|---|
| Demographic information: | | |
| Number of participants | 29(14 females) | |
| **Age**(yrs) | 26.34 | 7.09 |
| Questionnaires: | | |
| QIDS | 8.59 | 5.23 |
| STAI | 86.00 | 20.35 |
| TEPS | 77.34 | 11.83 |
| RRS | 47.00 | 13.61 |

Note: QIDS: the Quick Inventory of Depressive Symptoms. STAI: Spielberger State-Trait Anxiety Inventory. TEPS: the Temporal Experience of Pleasure Scale. RRS: the Ruminative Response Scale. SD: standard deviation.

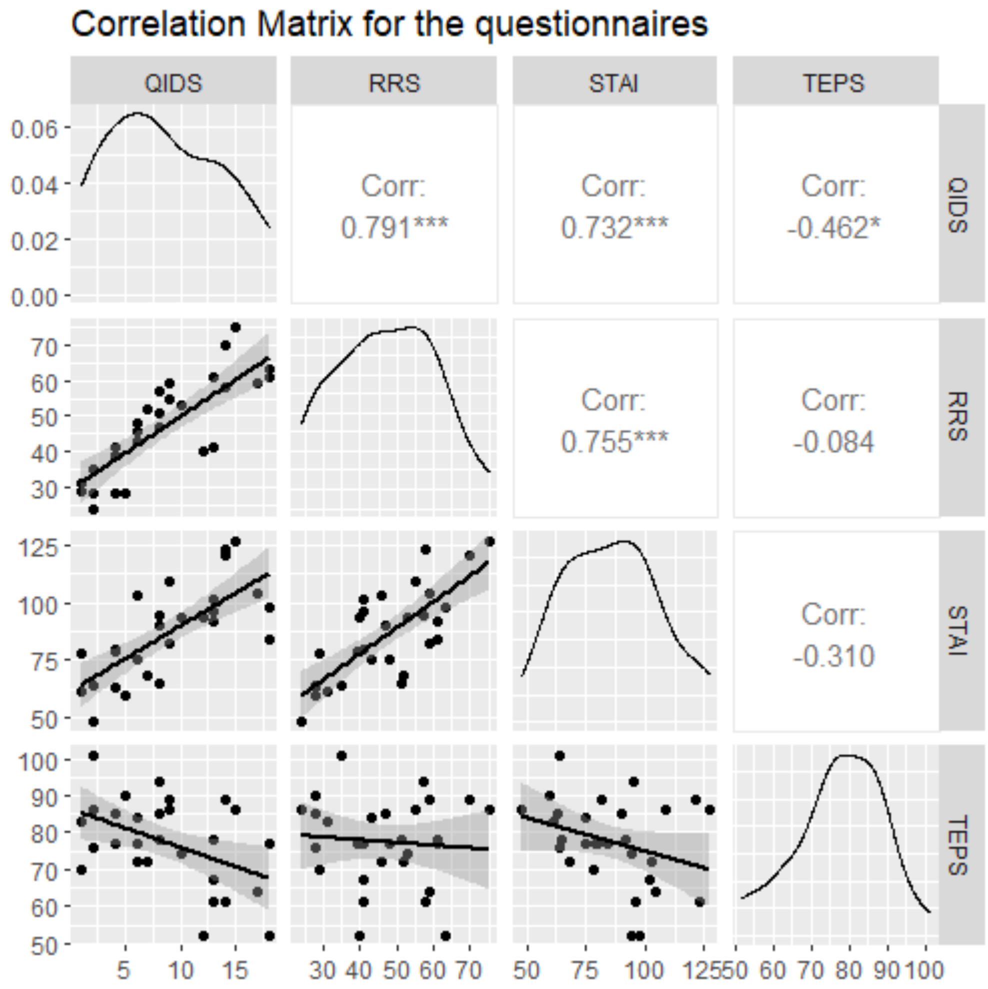


**Fig.S 1 |** Correlation Matrix for the questionnaires.

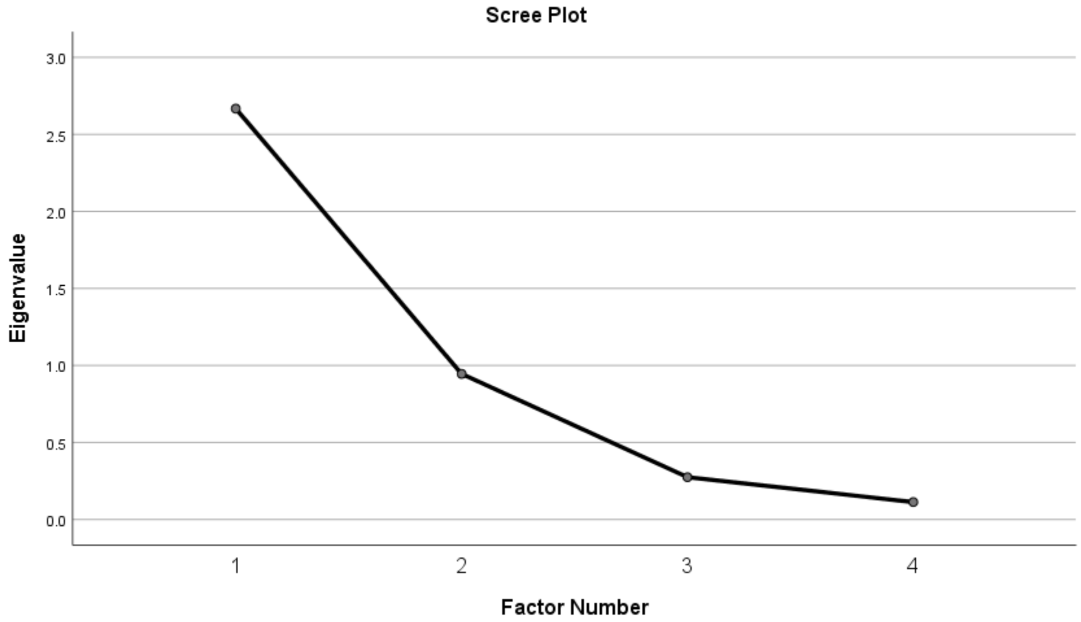


**Fig.S 2 | the Scree Plot of the factor analysis.**

There was no significant correlation between the overall depression scores and any of the base learning rates (all p>.656).

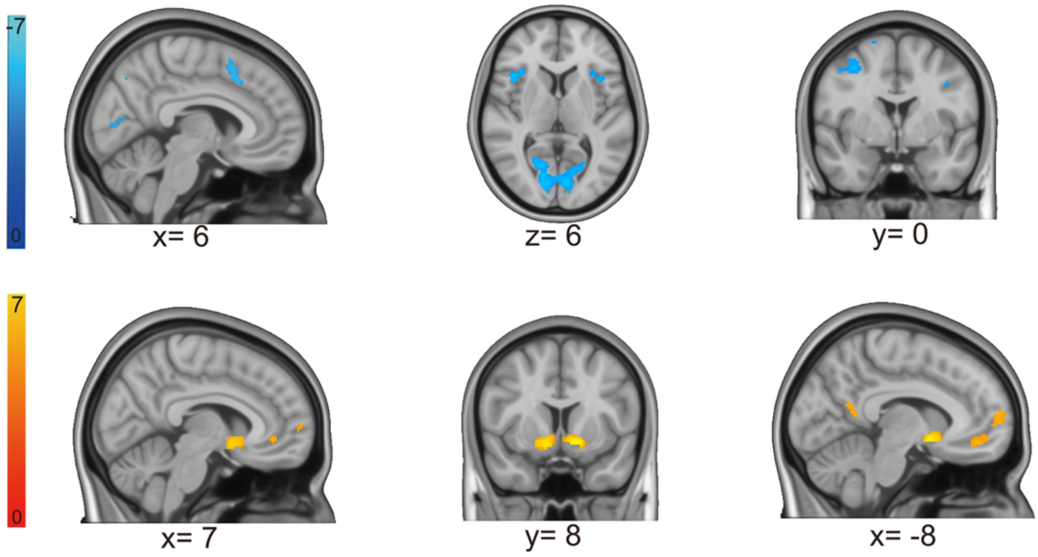


**Fig.S 3| brain regions obtained for the positive (win + no loss) minus negative outcomes contrast, i.e., win and no-loss minus no-win and loss (z>4.6, cluster-wise P<0.05).**

#### ANCOVA results for choice accuracies

We examined the influences of an informativity imbalance formally by running a valence (win vs loss)*self-volatility (volatile vs stable)*other-volatility (volatile vs stable)*negativity scores ANCOVA analysis, with self-volatility being the volatility level for the specific outcome (e.g. self-volatility is volatile for win accuracies in the win-volatile block) and other-volatility as the volatility level for outcomes of the other valance in the same block ( e.g. in the win-volatile block, for win accuracy, self-volatility (i.e. win) is volatile and other-volatility (i.e. loss) is stable). For example, for wins, the both volatile and the win-volatile block were self(win)-volatile, whereas the loss-volatile block and the both-stable block were self(win)-stable.), and other-volatility ("other volatile"/"other-stable", e.g. for wins, the both volatile and the loss-volatile block were other(loss)-volatile, whereas the win-volatile block and the both-stable block were other(loss)-stable.) as repeated measures.

This analysis confirmed a significant main effect of self-volatility ($F(1,27)=35.839$, $p<.001$, $\eta_p^2=.570$), showing that volatility increased accuracies in general, as well as a significant main effect of other-volatility i.e., the volatility levels of the competing information ($F(1,27)=105.193$, $p<.001$, $\eta_p^2=.796$), showing that accuracy decreased when the competing information was more volatile. Importantly, the analysis revealed a significant interaction effect of self-volatility * other-volatility ($F(1,27)=112.360$, $p<.001$, $\eta_p^2=.806$). This result suggests, as we showed in the main text, when there was an imbalance in informativity of the information (i.e., one being more volatile than the other), participants attended and learned more from the more volatile/informative feedback. Interestingly, we also found a significant self-volatility * other-volatility*negativity scores interaction ($F(1,27)=4.448$, $p=.044$, $\eta_p^2=.141$).

We did not find a significant main effect of valence (p=0.245), however, we found a marginally significant valance*self-volatility*negativity scores interaction ($F(1,27)=3.579$, $p=.069$, $\eta_p^2=.117$), showing higher self-volatility effects for win relative to loss for people with higher negativity scores.

Table. S 2 valence * self-volatility*other-volatility negativity scores ANCOVA results for choice accuracies

| Source | Type III Sum of Squares | df | Mean Square | F | Sig. | Partial Eta Squared |
|---|---|---|---|---|---|---|
| valence | .007 | 1 | .007 | 1.413 | .245 | .050 |
| valence * negativity_scores | .002 | 1 | .002 | .466 | .501 | .017 |
| Error(valence) | .129 | 27 | .005 | | | |
| self_volatility | .087 | 1 | .087 | 35.839 | <.001 | .570 |
| self_volatility * negativity_scores | .000 | 1 | .000 | .077 | .784 | .003 |
| Error(self_volatility) | .066 | 27 | .002 | | | |
| other_volatility | .317 | 1 | .317 | 105.193 | <.001 | .796 |
| other_volatility * negativity_scores | .004 | 1 | .004 | 1.191 | .285 | .042 |
| Error(other_volatility) | .081 | 27 | .003 | | | |
| valence * self_volatility | .017 | 1 | .017 | 1.299 | .264 | .046 |
| valence * self_volatility * negativity_scores | .047 | 1 | .047 | 3.579 | .069 | .117 |
| Error(valence*self_volatility) | .358 | 27 | .013 | | | |
| valence * other_volatility | .001 | 1 | .001 | .059 | .810 | .002 |
| valence * other_volatility * negativity_scores | .011 | 1 | .011 | .718 | .404 | .026 |
| Error(valence*other_volatility) | .421 | 27 | .016 | | | |
| self_volatility * other_volatility | .334 | 1 | .334 | 112.360 | <.001 | .806 |
| self_volatility * other_volatility * negativity_scores | .013 | 1 | .013 | 4.448 | .044 | .141 |
| Error(self_volatility*other_volatility) | .080 | 27 | .003 | | | |
| valence * self_volatility * other_volatility | .001 | 1 | .001 | .380 | .543 | .014 |
| valence * self_volatility * other_volatility * negativity_scores | 4.325E-6 | 1 | 4.325E-6 | .001 | .973 | .000 |
| Error(valence*self_volatility*other_volatility) | .097 | 27 | .004 | | | |

#### ANCOVA results for model estimates.

Table. S 3 valence * self-volatility*other-volatility negativity scores ANCOVA results for learning rates (alpha)

| Source | Type III Sum of Squares | df | Mean Square | F | Sig. | Partial Eta Squared |
|---|---|---|---|---|---|---|
| valence | 16.897 | 1 | 16.897 | 4.578 | .042 | .145 |
| valence * negativity_scores | 1.828 | 1 | 1.828 | .495 | .488 | .018 |
| Error(valence) | 99.652 | 27 | 3.691 | | | |
| self_volatility | 17.634 | 1 | 17.634 | 20.444 | <.001 | .431 |
| self_volatility * negativity_scores | .077 | 1 | .077 | .089 | .767 | .003 |
| Error(self_volatility) | 23.289 | 27 | .863 | | | |
| other_volatility | .815 | 1 | .815 | .284 | .598 | .010 |
| other_volatility * negativity_scores | 1.744 | 1 | 1.744 | .609 | .442 | .022 |
| Error(other_volatility) | 77.372 | 27 | 2.866 | | | |
| valence * self_volatility | .059 | 1 | .059 | .074 | .787 | .003 |
| valence * self_volatility * negativity_scores | 4.498 | 1 | 4.498 | 5.668 | .025 | .173 |
| Error(valence*self_volatility) | 21.430 | 27 | .794 | | | |
| valence * other_volatility | 5.461 | 1 | 5.461 | 2.248 | .145 | .077 |
| valence * other_volatility * negativity_scores | 1.557 | 1 | 1.557 | .641 | .430 | .023 |
| Error(valence*other_volatility) | 65.582 | 27 | 2.429 | | | |
| self_volatility * other_volatility | 8.792 | 1 | 8.792 | 11.135 | .002 | .292 |
| self_volatility * other_volatility * negativity_scores | 1.328 | 1 | 1.328 | 1.682 | .206 | .059 |
| Error(self_volatility*other_volatility) | 21.318 | 27 | .790 | | | |
| valence * self_volatility * other_volatility | 1.729 | 1 | 1.729 | 2.912 | .099 | .097 |
| valence * self_volatility * other_volatility * negativity_scores | .662 | 1 | .662 | 1.116 | .300 | .040 |
| Error(valence*self_volatility*other_volatility) | 16.034 | 27 | .594 | | | |

Our winning model has an inverse temperature term per block. Here we run a valence * volatility * negativity scores ANCOVA, with valence being win or loss, and volatility being volatile vs stable, so valence*volatility corresponds to all four block types.

**Table. S 4 valence * volatility* negativity scores ANCOVA results for inverse temperatures (beta)**

| Source | Type III Sum of Squares | df | Mean Square | F | Sig. | Partial Eta Squared |
|---|---|---|---|---|---|---|
| valence | .009 | 1 | .009 | .032 | .860 | .001 |
| valence * negativity_scores | 1.249 | 1 | 1.249 | 4.396 | .046 | .140 |
| Error(valence) | 7.673 | 27 | .284 | | | |
| volatility | .898 | 1 | .898 | 4.254 | .049 | .136 |
| volatility * negativity_scores | .001 | 1 | .001 | .006 | .939 | .000 |
| Error(volatility) | 5.702 | 27 | .211 | | | |
| valence * volatility | 5.225 | 1 | 5.225 | 34.812 | <.001 | .563 |
| valence * volatility * negativity_scores | .079 | 1 | .079 | .529 | .473 | .019 |
| Error(valence*volatility) | 4.053 | 27 | .150 | | | |

#### ANCOVA results for fMRI ROIs.

**Table. S 5 region (pgACC vs. dACC) * valence * self-volatility* negativity scores ANCOVA results for ROI results**

| Source | Type III Sum of Squares | df | Mean Square | F | Sig. | Partial Eta Squared |
|---|---|---|---|---|---|---|
| region | 5.852 | 1 | 5.852 | 10.037 | .004 | .271 |
| region * negativity_scores | 3.033 | 1 | 3.033 | 5.201 | .031 | .162 |
| Error(region) | 15.743 | 27 | .583 | | | |
| valence | 2.075 | 1 | 2.075 | 2.130 | .156 | .073 |
| valence * negativity_scores | .405 | 1 | .405 | .415 | .525 | .015 |
| Error(valence) | 26.307 | 27 | .974 | | | |
| volatility | .994 | 1 | .994 | 1.911 | .178 | .066 |
| volatility * negativity_scores | 3.806 | 1 | 3.806 | 7.316 | .012 | .213 |
| Error(volatility) | 14.048 | 27 | .520 | | | |
| region * valence | 83.883 | 1 | 83.883 | 173.122 | <.001 | .865 |

| | | | | | | |
|---|---|---|---|---|---|---|
| region * valence * negativity_scores | .041 | 1 | .041 | .084 | .775 | .003 |
| Error(region*valence) | 13.082 | 27 | .485 | | | |
| region * volatility | .008 | 1 | .008 | .018 | .893 | .001 |
| region * volatility * negativity_scores | .005 | 1 | .005 | .010 | .921 | .000 |
| Error(region*volatility) | 12.324 | 27 | .456 | | | |
| valence * volatility | .124 | 1 | .124 | .198 | .660 | .007 |
| valence * volatility * negativity_scores | .026 | 1 | .026 | .042 | .840 | .002 |
| Error(valence*volatility) | 16.857 | 27 | .624 | | | |
| region * valence * volatility | 5.378 | 1 | 5.378 | 20.566 | <.001 | .432 |
| region * valence * volatility * negativity_scores | 1.827 | 1 | 1.827 | 6.985 | .014 | .206 |
| Error(region*valence*volatility) | 7.061 | 27 | .262 | | | |

### References


Behrens, T. E., Woolrich, M. W., Walton, M. E., & Rushworth, M. F. (2007). Learning the value of information in an uncertain world. *Nature neuroscience*, *10*(9), 1214-1221.

Bowling, S. R., Khasawneh, M. T., Kaewkuekool, S., & Cho, B. R. (2009). A logistic approximation to the cumulative normal distribution. *Journal of industrial engineering and management*, *2*(1), 114-127.

Brooks, J. C., Beckmann, C. F., Miller, K. L., Wise, R. G., Porro, C. A., Tracey, I., & Jenkinson, M. (2008). Physiological noise modelling for spinal functional magnetic resonance imaging studies. *Neuroimage*, *39*(2), 680-692.

Brooks, S. P., & Gelman, A. (1998). General methods for monitoring convergence of iterative simulations. *Journal of computational and graphical statistics*, *7*(4), 434-455.

Carpenter, B., Gelman, A., Hoffman, M. D., Lee, D., Goodrich, B., Betancourt, M., Brubaker, M., Guo, J., Li, P., & Riddell, A. (2017). Stan: A probabilistic programming language. *Journal of statistical software*, *76*(1).

Gard, D. E., Gard, M. G., Kring, A. M., & John, O. P. (2006). Anticipatory and consummatory components of the experience of pleasure: A scale development study. *Journal of Research in Personality*, *40*(6), 1086-1102. https://doi.org/https://doi.org/10.1016/j.jrp.2005.11.001

Jenkinson, M. (2004). Improving the registration of B0-disorted Epi images using calculated cost function weights: we 202. *Neuroimage*, *22*.

Jenkinson, M., Bannister, P., Brady, M., & Smith, S. (2002). Improved optimization for the robust and accurate linear registration and motion correction of brain images. *Neuroimage*, *17*(2), 825-841.

Jenkinson, M., Beckmann, C. F., Behrens, T. E., Woolrich, M. W., & Smith, S. M. (2012). Fsl. *Neuroimage*, *62*(2), 782-790.

Nolen-Hoeksema, S., & Morrow, J. (1991). A prospective study of depression and posttraumatic stress symptoms after a natural disaster: the 1989 Loma Prieta Earthquake. *Journal of personality and social psychology*, *61*(1), 115.

Pulcu, E., & Browning, M. (2017). Affective bias as a rational response to the statistics of rewards and punishments. *Elife*, *6*, e27879.

Rescorla, R. A. (1972). A theory of Pavlovian conditioning: Variations in the effectiveness of reinforcement and nonreinforcement. *Current research and theory*, 64-99.

Rush, A. J., Trivedi, M. H., Ibrahim, H. M., Carmody, T. J., Arnow, B., Klein, D. N., Markowitz, J. C., Ninan, P. T., Kornstein, S., & Manber, R. (2003). The 16-Item Quick Inventory of Depressive Symptomatology (QIDS), clinician rating (QIDS-C), and self-report (QIDS-SR): a psychometric evaluation in patients with chronic major depression. *Biological psychiatry*, *54*(5), 573-583.

Shrout, P. E., & Bolger, N. (2002). Mediation in experimental and nonexperimental studies: new procedures and recommendations. *Psychological methods*, *7*(4), 422.

Silvetti, M., Seurinck, R., & Verguts, T. (2013). Value and prediction error estimation account for volatility effects in ACC: A model-based fMRI study. *Cortex*, *49*(6), 1627-1635. https://doi.org/https://doi.org/10.1016/j.cortex.2012.05.008

Smith, S. M. (2002). Fast robust automated brain extraction. *Human brain mapping*, *17*(3), 143-155.

Spielberger, C. D. (1983). State-trait anxiety inventory for adults.

Vehtari, A., Gelman, A., & Gabry, J. (2017). Practical Bayesian model evaluation using leave-one-out cross-validation and WAIC. *Statistics and Computing*, *27*(5), 1413-1432. https://doi.org/10.1007/s11222-016-9696-4
